# PGC1α and Exercise Adaptations in Zebrafish

**DOI:** 10.1101/483784

**Authors:** Alice Parisi, Peter Blattmann, Giulia Lizzo, Vivienne Stutz, Laura Strohm, Joy Richard, Gabriele Civiletto, Aline Charpagne, Frederic Raymond, Cedric Gobet, Benjamin Weger, Eugenia Migliavacca, Ruedi Aebersold, Bruce Spiegelman, Philipp Gut

## Abstract

Fish species display huge differences in physical activity ranging from lethargy to migration of thousands of miles, making them an interesting model to identify determinants of physical fitness. Here, we show a remarkable plasticity of zebrafish in response to exercise and induction of PGC1α (encoded by *PPARGC1A*), a dominant regulator of mitochondrial biogenesis. Forced expression of human *PPARGC1A* induces mitochondrial biogenesis, an exercise-like gene expression signature, and physical fitness comparable to wild-type animals trained in counter-current swim tunnels. Quantifying transcriptional and proteomic changes in response to exercise or PGC1α, we identify conserved ‘exercise’ adaptations, including a stoichiometric induction of the electron transport chain (ETC) that re-organizes into respiratory supercomplexes in both conditions. We further show that ndufa4/ndufa4l, previously assigned to complex I, associates to free and supramolecular complex IV *in vivo*. Thus, zebrafish is a useful and experimentally tractable vertebrate model to study exercise biology, including ETC expression and assembly.

**HIGHLIGHTS:**

- PGC1α reprograms zebrafish skeletal muscle to a ‘red fiber’ phenotype and increases exercise performance
- Zebrafish show a high molecular plasticity in response to PGC1α and exercise
- SWATH-MS proteomics show a stoichiometric induction of the electron transport chain that organizes as supercomplexes in response to PGC1α and exercise
- ndufa4/ndufa4l associate to free and supramolecular complex IV *in vivo*

## INTRODUCTION

Some fish species, in particular tuna and salmonids, are considered extraordinary athletes based on their biomechanical and cardio-metabolic properties that allow them to sustain high swimming performances over extended periods of time. This includes migration from spawning to distant feeding habitats (Block et al., 2005). Zebrafish, a commonly used laboratory animal, show low levels of activity in their natural habitat, but are capable of remarkable physical performance when trained in counter-current swim tunnels; chronic endurance training has been reported to activate angiogenic and myogenic programs and to increase aerobic capacity (Gilbert et al., 2014; Palstra et al., 2014). This plasticity suggests that zebrafish can be used as an informative vertebrate model to understand the underlying mechanisms by which cells and tissues adapt in response to physical challenges, such as physiological adaptations in response to exercise training in humans. In particular, packs of zebrafish can be exposed, easily and inexpensively, to strenuous physical challenges over extended periods of time. The use of zebrafish for exercise biology has been limited in the past, in part due to a lack of understanding of the underlying genetic pathways in fish. Establishing exercise paradigms in zebrafish will facilitate the dissection of the pathways that control physical plasticity through forward- and reverse genetics technologies as well as by applying reporter tools to monitor physiological processes *in vivo* (Gut et al., 2017; Lieschke and Currie, 2007).

Research in the past twenty years has identified peroxisome proliferator-activated receptor γ coactivator 1-alpha (PGC1α, encoded by the *Ppargc1a* gene) as a master regulator of cell intrinsic and extrinsic exercise adaptations, including mitochondrial biogenesis and function, neovascularization, formation of neuromuscular junctions, and switching from fast to slow-twitch muscle fibers in humans and mice (Handschin and Spiegelman, 2008). Muscle-specific transgenic expression of *Ppargc1a* in mice leads to a macroscopic red appearance of skeletal muscle due to the high content of oxidative slow-twitch muscle fibers, and increases resistance to fatigue of myofibers *ex vivo* (Lin et al., 2002). More recently, proteomics and metabolomics profiling strategies have been used in mice to identify secreted peptides and metabolites that have systemic effects on energy metabolism (Bostrom et al., 2012; Rao et al., 2014; Roberts et al., 2014), cognitive functions (Wrann et al., 2013), and mood (Agudelo et al., 2014).

In this study we test molecular and physiological adaptations of zebrafish by testing the evolutionary conservation of downstream effects that have been attributed to PGC1α activity in mammals, and subsequently compare the effects of PGC1α to that of chronic endurance training. We show that forced expression of human *PPARGC1a* is sufficient to induce macroscopic changes of skeletal muscle that are reminiscent to the myoglobin-rich, aerobic red fiber appearance seen in transgenic *PPARGC1A* mice. Developing an exercise protocol in counter-current swim tunnels we profile the effects of exercise or PGC1α on a molecular level. To this end, we carry out transcriptome profiling using RNA-Seq and a precise proteomic profiling via sequential window acquisition of all theoretical spectral (SWATH) mass spectrometry, a technique that is ideally suited for comparative, quantitative analyses of complex proteomes (Gillet et al., 2012). The data show a high evolutionary conservation of exercise adaptations and PGC1α effects at the both transcriptome and proteome level. Among those, we find a stoichiometric induction of the electron transport chain (ETC); we leverage this dataset of ‘molecular exercise adaptation’ to confirm ndufa4l, the orthologue to the atypical ETC subunit NDUFA4, as a subunit primarily associating to complex IV (CIV) (Balsa et al., 2012). Furthermore, we find unexpectedly that the ETC components of zebrafish becomes rapidly stored into respiratory supercomplexes in response to short-term exercise. These results highlight the potential of zebrafish to study evolutionary conserved pathways that regulate cellular responses to chronic exercise and indicate how PGC1α and chronic exercise affect the reorganization of the electron transport chain.

Thus, we report the first example of a genetic manipulation in zebrafish that increases physical performance and induces adaptations reminiscent of responses that occur in trained mammals setting the stage for the genetic deconvolution of the exercise/ PGC1α program in a vertebrate.

## RESULTS

### Transgenic expression of human PGC1α in skeletal muscle of zebrafish

Functional domains of PGC1α protein are largely conserved among zebrafish, mouse and human (Figure S1). We generated stable transgenic zebrafish lines that express an additional copy of zebrafish *ppargc1a* or human *PPARGC1A* using a muscle-specific promoter: *Tg(actcb1:ppargc1a;cryaa:ZsGreen1)^nei1^* and *Tg(actcb1:PPARGC1A;cryaa:ZsGreen1) ^nei2^* (hereafter termed *actc1b:*PGC1α) (Figure 1A). Whereas the transgenic zebrafish larvae are macroscopically indistinguishable from wild-type siblings (data not shown), adult transgenic carriers are characterized by a red body appearance in early adulthood starting at 6 weeks post fertilization (Figure S2). The trunk skeletal muscle shows a ‘red meat’ phenotype reminiscent to what has been observed in transgenic mice overexpressing murine PGC1α (Figure 1B and 1C) (Lin et al., 2002). This phenotype observed in the transgenic line carrying human *PPARGC1A* suggests a high evolutionary conservation since the human PGC1α protein must be structurally capable to interact with endogenous transcription factors that mediate PGC1α signals (Lin et al., 2005). As we observed identical phenotypes when overexpressing the zebrafish ppargc1a, (data not shown), we selected the transgenic line expressing human PGC1α to study its molecular gain-of-function phenotypes. The *actc1b:*PGC1α transgene expression was approximately 25-fold higher relative to the endogenous *ppargc1a* transcript at four months of age (Figure 1D). Since the red color of muscle is influenced by a high myoglobin (encoded by *mb*) content, we quantified *mb* expression levels and found an eight-fold increase in *actc1b:*PGC1α zebrafish compared to control animals (Figure 1E). Western blotting for porin, a marker for mitochondrial content, showed an increase in mitochondrial biogenesis (Figure 1F), consistent with an increase of citrate synthase and respiratory chain complex activities in *actc1b:*PGC1α animals (Figure S3). To quantitatively assess the transcriptional changes induced by PGC1α, we performed microarray analysis comparing control and *actc1b:*PGC1α adult skeletal muscles. To precisely analyze the effects of PGC1α on the expression profile of exercise-related genes, we performed an enrichment analysis on a literature- and gene-ontology based list of marker genes of pathways that are known to be regulated by chronic exercise. These pathways include genes related to vascularization, neuromuscular junction formation and function, slow- and fast-twitch fiber identity, muscle development, mitochondria, and energy metabolism. Strikingly, a near complete suppression of the fast-twitch fiber program was detected in contrast to an up-regulation of a slow-twitch fiber signature. This is paralleled by an increased expression of slow-twitch fiber genes (Figure 1G). Consistent with a shift towards oxidative myofibers, we found that mitochondrial biogenesis and metabolic genes were upregulated. Three quartiles of all genes associated with the mitochondria gene set were induced larger than 2-fold.

**Figure 1.**
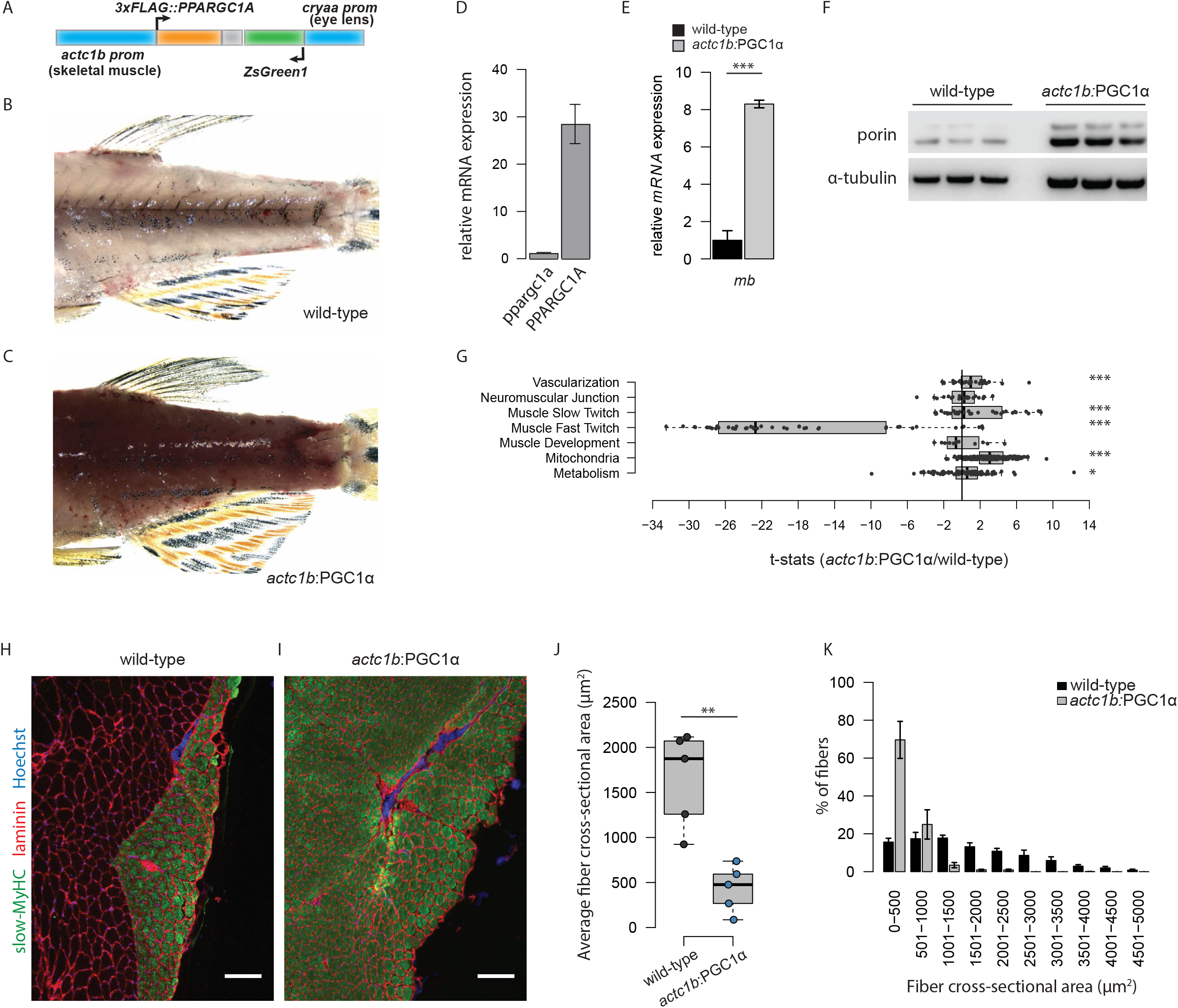
Human PGC1α reprograms energy metabolism and induces fiber type switching. (A) Transgenic construct used for overexpression of *PPARGC1A*. An eye-marker cassette expressing *ZsGreen1* was used to identify transgenic animals as early as 48hpf. (B and C) Lateral view of trunks of a wild-type (B) and *actc1b:*PGC1α (C) zebrafish at 4 months of age. (D) Relative *mRNA* expression of human *PPARGC1A* relative to endogenous *ppargc1a* in *actc1b:*PGC1α adults (n=4). (E) Relative *mRNA* expression of *myoglobin* relative to *bactin*, in wild-type and *actc1b:*PGC1α adult fish (n=4 per genotype). (F) Western blot analysis of the mitochondrial marker protein porin in wild-type and *actc1b:*PGC1α adult fish (n=3 per genotype). (G) Enrichment test of known exercise-regulated genes in skeletal muscle of *actc1b:*PGC1α zebrafish compared to wild-type animals as measured by customized gene expression arrays (n=4 per genotype). Statistics represented correspond to p-adjusted values, calculated by Benjamini-Hochberg (HB) multiple test. (H and I) Immunofluorescence staining of slow-myosin heavy chain as marker for slow-twitch myofibers and laminin to mark the basal lamina of myofibers in trunk muscle of wild-type (H) and *actc1b:*PGC1α zebrafish (I) (Scale bar, 100 μm). (J) Quantification of average cross-sectional area of trunk muscle fibers in wild-type and transgenic animals (n=5 per genotype; average counts from at least 250 myofibers per animal). (K) Frequency distribution of cross-sectional fiber area in wild-type and PGC1α zebrafish (n=5 per genotype; counts from at least 250 myofibers per animal). p = 0.0233, Kolmogorov Smirnov (KS) test. All measurements and images analyzed at 4 months of age. *mb* = *myoglobin*. * p < 0.05, ** p < 0.01, *** p < 0.001.

Anatomically, fiber type distribution in zebrafish, similar to most fish species, is organized by a thin layer of slow-twitch fibers along the flanks for the regulation of posture, while the majority of the trunk consists of fast-twitch or intermediate fiber types for swimming (Altringham and Ellerby, 1999). Immunofluorescent staining of transversal sections revealed that *actc1b:*PGC1α trunk muscle exclusively consists of slow muscle fibers (Figure 1H and 1I). Consistent with this finding, the average fiber size was strongly reduced with a drastic shift to fibers of a cross-sectional area smaller than 500μm^2^ (Figure 1J and 1K). These results indicate that the human PGC1α protein is sufficient to induce in zebrafish anatomic and molecular features of exercise adaptations in an evolutionary conserved manner.

### PGC1α increases physical performance without prior training

We next established an aerobic training protocol using counter-current swim tunnels to compare the exercise performance of untrained *actc1b:*PGC1α zebrafish with that of wild-type animals. Of note, whereas rodents need to be forced by strong stimuli (such as shock grid, tail tapping, or high pressure bursts of air) to exercise (Knab et al., 2009), zebrafish voluntarily and consistently swim against the water current. To this end, a pack of 8 zebrafish is placed into an immersed chamber that is exposed to defined currents (Movie S1).

After determining the baseline critical swimming speed (Ucrit) (Gilbert et al., 2014) of wild-type zebrafish using an endurance test, we trained zebrafish by setting an initial current of 50 cm/s (which corresponds to 75% of their initial Ucrit) for 3 days followed by 4 weeks of training that included exercising the zebrafish for 3 hours on 5 days per week using incremental currents (Figure 2A). Control groups of untrained *actc1b:*PGC1α or wild-type zebrafish were placed in a chamber at a baseline current of 5 cm/s. An endurance test was performed during week 4 using an individual swim tunnel, and animals were sacrificed at the end of the week. Four weeks of exercise were sufficient to induce a significant increase of the UCrit in wild-type animals. Strikingly, the Ucrit values of untrained transgenic animals were similar to that of the wild-type training group without the need for prior exercise training (Figure 2B). Consistent with exercise adaptations, increased tomm20 expression indicated that mitochondrial biogenesis was induced in the training group, albeit to a lesser extent than in *actc1b:*PGC1α animals (Figure 2C and 2D).

**Figure 2.**
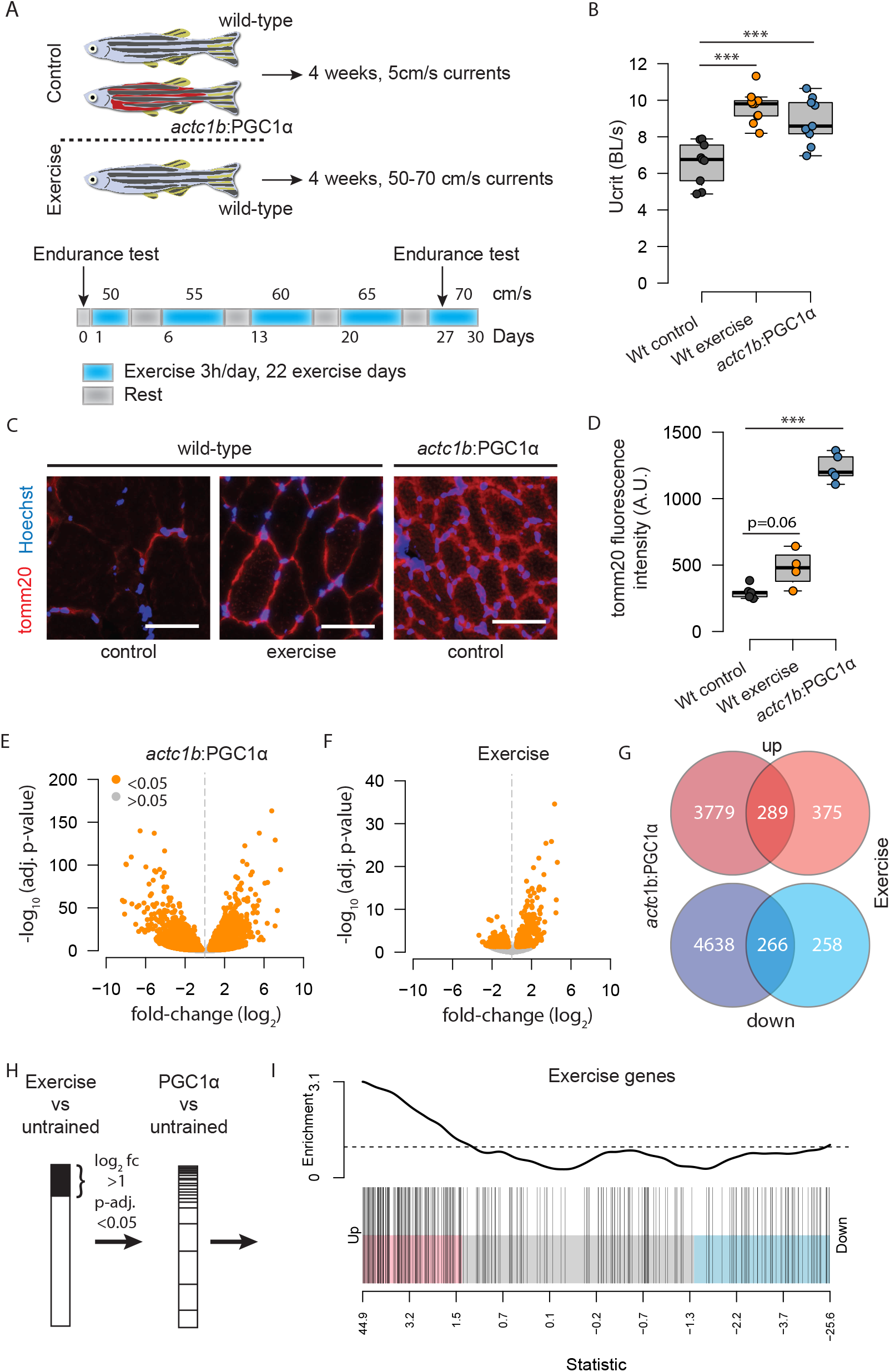
PGC1α increases physical performance and induces an ‘exercise’ gene signature. (A) Schematic representation of exercise protocol. (B) Critical swimming speed (Ucrit) as measured through an endurance test in the final week of the exercise intervention. (C) Immunofluorescence staining of the mitochondrial marker protein tomm20 in each group and (D) quantification of fluorescence intensity (n=4 per group; average count of at least 250 myofibers per animal) (Scale bar: 50 μm). Muscles were collected from animals subjected to an independent but identical exercise program after three weeks. (E) Volcanoplot of relative gene expression in *actc1b:*PGC1α and (F) exercised zebrafish compared to the wild-type control animals (n=8 per group). Differentially regulated genes in orange (log2 fold-change > 0.5, p-adj. < 0.05). (G) Venn diagram of differentially regulated genes. (H) Barcode plot of an ‘exercise-responsive gene set’ shows enrichment of the same gene set in the transcriptome of *actc1b:*PGC1α animals compared to controls. WT: wild-type, A.U.: arbitrary units. *** p < 0.001

### PGC1α induces an exercise-like gene signature

Trunk muscle tissues of the control, exercised and *actc1b:*PGC1α animals were dissected for RNA extraction and RNA-seq analysis. Consistent with the complete remodeling of the skeletal muscle observed in *actc1b:*PGC1α animals, we found a major transcriptomic reprogramming with 4068 genes up-regulated and 4904 genes down-regulated upon PGC1α overexpression compared to untrained control animals (log2 fold-change >1; p-adj. <0.05, n=8. Figure 2E and 2G). In contrast, 4 weeks of daily exercise of wild-type animals had a less pronounced effect on gene expression, with 644 genes up- and 524 genes being down-regulated (Figure 2F and 2G). The drastic reprogramming of approximately 20% of the transcriptome in transgenic animals compared to less than 2% in response to exercise as a physiological stimulus suggests that PGC1α overexpression has a potent effect on transcriptome activation with potentially artificially forced expression on a subset of genes. Indeed, we observed increased expression of several genes that are not expressed in skeletal muscle under wild-type conditions (data not shown). We therefore defined the ‘exercise gene signature’ as the genes significantly regulated in exercised fish compared to wild-type controls (log2 fold-change >1; p-adj. <0.05, n=8, 1168 genes. Table S2). Among these genes, we found many genes linked to muscle physiology that are known to be regulated by exercise and PGC1α: for example *peroxisome proliferator activated receptor α* (*ppara*) and *estrogen-related receptor alpha* (*essra*), two well-established interactors of PGC1α regulating mitochondrial metabolism were significantly induced (Lin et al., 2005; Mootha et al., 2004; Vega et al., 2000). However, we also identified genes that are known to be critical for skeletal muscle physiology, but that have not been linked previously to PGC1α activation. These include *nicotinamide riboside kinase 2* (*nmrk2*) (Fletcher et al., 2017), the rate-limiting enzyme of NAD^+^ generation from Vitamin B3 precursors, *lim homeobox domain 2a* (*lhx2a*) (Kodaka et al., 2015), a repressor of myotube differentiation, or *myomaker* (*mymk*), a small peptide regulating myotube fusion (Goh and Millay, 2017; Millay et al., 2013) (Table 1).

**Table 1.** Examples of genes differentially expressed after exercise and by PGC1α overexpression.

| Name | log<sub>2</sub>FC<br>Exercise | p-adj. | log<sub>2</sub>FC<br>PGC1α | p-adj. | Encoded protein \| function* | |
| --- | --- | --- | --- | --- | --- | --- |
| ***Energy metabolism and mitochondria*** |  |  |  |  |  |  |
| *nr4a3* | 4.61 | 2.78E-25 | 0.44 | 0.576 | NOR1 \| Exercise-induced transcription factor drives muscle reprogramming, vascularization, oxidative metabolism and endurance | Goode et al. (2016) |
| *nmrk2* | 2.72 | 1.78E-15 | 1.63 | 4.37E-07 | nrk2 \| Rate-limiting enzyme of the NAD<sup>+</sup> synthesis pathway | (Fletcher et al., 2017) |
| *esrra* | 2.01 | 1.06E-05 | 0.84 | 0.0607 | errα \| Transcription factor regulating mitochondrial biogenesis and oxidative metabolism | (Schreiber et al., 2004) |
| *klf15* | 1.34 | 0.0008 | -0.097 | 0.0052 | klf15 \| Exercise-induced transcription factor regulating lipid oxidation | (Haldar et al., 2012) |
| *ucp3* | 1.085 | 0.038 | -0.23 | 0.6542 | ucp3 \| Muscle-specific mitochondrial uncoupling protein | (Boss et al., 1997) |
| *plin3* | 1.07 | 0.0006 | 1.98 | 2.47E-15 | perilipin 3 \| Lipid droplet-associated protein | (Covington et al., 2014) |
| *pparaa* | 1.05 | 0.0024 | 0.83 | 0.0041 | pparα \| Transcription factor regulating oxidative metabolism | (Ajith and Jayakumar, 2016) |
| ***Myogenesis and muscle regeneration*** |  |  |  |  |  |  |
| *mymk* | 2.98 | 0.0006 | 1.97 | 0.010217 | myomaker \| Peptide regulating fusion of adult myoblasts and that is required for muscle regeneration | (Millay et al., 2014) |
| *hgfa* | 2.87 | 1.04E-09 | 0.076 | 0.91 | hepatocyte growth factor \| Regulates muscle stem cell activation | (Anderson, 2016) |
| *cited4b* | 2.79 | 1.88E-08 | 0.98 | 0.055 | cited4 Stimulates cardiac muscle hypertrophy and regeneration after cardiac ischemic damage | (Bezzarides et al., 2016) |
| *trim63a* | 1.89 | 8.27E-07 | -1.71 | 8.04E-07 | atrogen-1 \| Ubiquitin ligase regulation protein turnover | (Gomes et al., 2001) |
| *klf4* | 1.51 | 0.02 | -1.13 | 0.0354 | klf4 \| Transcription factor regulating muscle cell fusion and metabolism | (Sunadome et al., 2011) |
| *myf5* | 1.47 | 0.0036 | 0.04 | 0.953 | myf5 \| transcription factor regulating myogenesis and renewal of myogenic precursor cells | (Zammit, 2017) |
| *myog* | -1.46 | 0.0196 | 0.09 | 0.894 | myogenin \| Transcription factor regulating differentiation of myoblasts to myotubes | (Zammit, 2017) |
| *pitx3* | -1.49 | 5.50E-09 | 1.21 | 3.50E-07 | pitx3 \| Transcription factor regulating fetal myogenesis, anti-oxidant defense, and mitochondrial biogenesis | (L'Honore et al., 2014) |
| *lhx2a* | -2.56 | 0.009883 | -4.59 | 5.05E-09 | lhx2a \| Transcription factor negatively regulating skeletal muscle differentiation | (Kodaka et al., 2015) |
| **Anti-oxidant defense** |  |  |  |  |  |  |
| *mafK* | 2.33 | 9.17E-07 | -1.857 | 9.35E-05 | mafK \| Transcription factor forming heterodimers with leucine zipper transcription factors, including nuclear respiratory factors | (Zhu and Fahl, 2001) |
| *nfe2l1a* | 1.05 | 0.028 | -2.01 | 9.69E-09 | nrf1 \| nuclear factor erythroid-2-like 1, a transcription factor inducing genes required for cellular stress response | (Kim et al., 2016) |
| *nfe2l2a* | 0.74 | 0.0083 | 0.33 | 0.185 | nrf2 \| Nuclear factor erythroid-2-like 1, a transcription factor inducing genes required for anti-oxidant defense | (Merry and Ristow, 2016) |
| **Autophagy and lysosomal function** |  |  |  |  |  |  |
| *pqlc2* | 2.81 | 5.82E-16 | -0.76 | 0.047 | pqlc2 \| Cationic amino acid transporter of lysosomes regulating amino acid homeostasis | (Liu et al., 2012) |
| *fnip2* | 2.38 | 1.94E-07 | -0.81 | 0.0887 | fnip2 \| Folliculin interacting protein that regulates mTOR pathway | (Takagi et al., 2008) |
| *fnip1* | 1.49 | 2.68E-17 | -0.14 | 0.539 | fnip1 \| Folliculin interacting protein proposed as a negative regulator of AMPK | (Reyes et al., 2015; Siggs et al., 2016) |
| *ulk2* | 1.46 | 3.40E-05 | 0.14 | 0.743 | ulk2 \| Protein kinase involved in autophagy initiation and glycolysis | (Li et al., 2016b) |
| *flcn* | 1.28 | 2.43E-05 | -0.622 | 0.0316 | folliculin \| scaffold protein involved in modulation of several cellular processes including the mTOR pathway, RhoA signaling, autophagy | (Schmidt and Linehan, 2018) |
| *nupr1* | 1.16 | 0.0049 | -2.04 | 6.27E-06 | nupr1 \| A chromatin-associated factor that modulates autophagy and metabolic stress response to glucose and oxygen deprivation | (Hamidi et al., 2013) |
| *tfe3b* | -0.94 | 0.019 | -3.07 | 6.93E-23 | tfe3b \| MITF family transcription factor regulating autophagy and lysosomal biogenesis | (Settembre et al., 2011) |
| **Angiogenesis** |  |  |  |  |  |  |
| *kdr1* | 0.99 | 0.009 | 1.08 | 0.000316 | vegfr2 \| tyrosine kinase (RTK) receptor for the vascular endothelial growth factor (vegfr) that mediate angiogenesis | (Shibuya, 2013) |
| *vegfab* | 0.53 | 0.103 | 2.06 | 1.17E-19 | vegfa \| vegfr1 and vegfr2 ligand, the main signal for proliferation, sprouting, migration and endothelial cell tube formation | (Ferrara et al., 2003) |
| *dre-mir-206-1* | -3.33 | 0.0001 | 1.49 | 0.015116 | Negative regulator of angiogenesis | (Stahlhut et al., 2012) |

We then rank-sorted the ‘exercise gene signature’ in the transcriptome of *actc1b:*PGC1α transgenic animals to calculate the distribution of this gene set across the entire transcriptome. Indeed, the exercise gene signature was significantly enriched in the transgenic animals, indicating that PGC1α overexpression induces transcriptional changes that are hallmarks of exercise even in the absence of training (p-adj< 10^-5^, n=8. Figure 2H and 2I). In summary, the transcriptional profiling showed that PGC1α overexpression was sufficient to induce expression of genes regulated by exercise.

### SWATH-MS quantification of zebrafish skeletal muscle proteome

As changes in gene expression do not necessarily translate into the protein level (Liu et al., 2016), we developed a SWATH-MS quantitative proteomic workflow to quantify the actual changes in protein expression in the zebrafish muscle (Röst et al., 2014; Schubert et al., 2015; Teo et al., 2015). To support the application of SWATH-MS in zebrafish, we generated a sample-specific spectral library supporting peptide centric analysis and extracted quantitative data for 885 proteins from the mass spectrometry-based acquisition of peptide intensities (Figure 3A and Table S3). Similar to what was observed on the transcriptomic level, protein expression levels were impacted more strongly by PGC1α overexpression than by exercise (Figure 3B and 3C). However, pathway analysis showed a consistent overlap in the pathways induced by PGC1a and exercise (Figure 3D, 3E, 3F and S4). Of note, the gene set ‘oxidative phosphorylation’ (OXPHOS) was ranked among the most prominently upregulated signatures induced both by exercise and PGC1α, both at the transcription and at the protein level. Remarkably, PGC1α potently induced large fold-changes and significant enrichment of almost the entire OXPHOS gene set. In contrast, exercise induced only modest fold-changes of the same gene set, but still nearly the entirety of the gene set was significantly upregulated (Figure S4). At the protein level, from the 129 proteins that showed increased expression levels in response to both exercise (log2 fold-change > 0.3, FDR < 0.01, n=8) and PGC1α overexpression (log2FC > 1, FDR < 0.01, n=8), 45 (34%) belong to the ETC (Figure 3D, Table S3).

**Figure 3.**
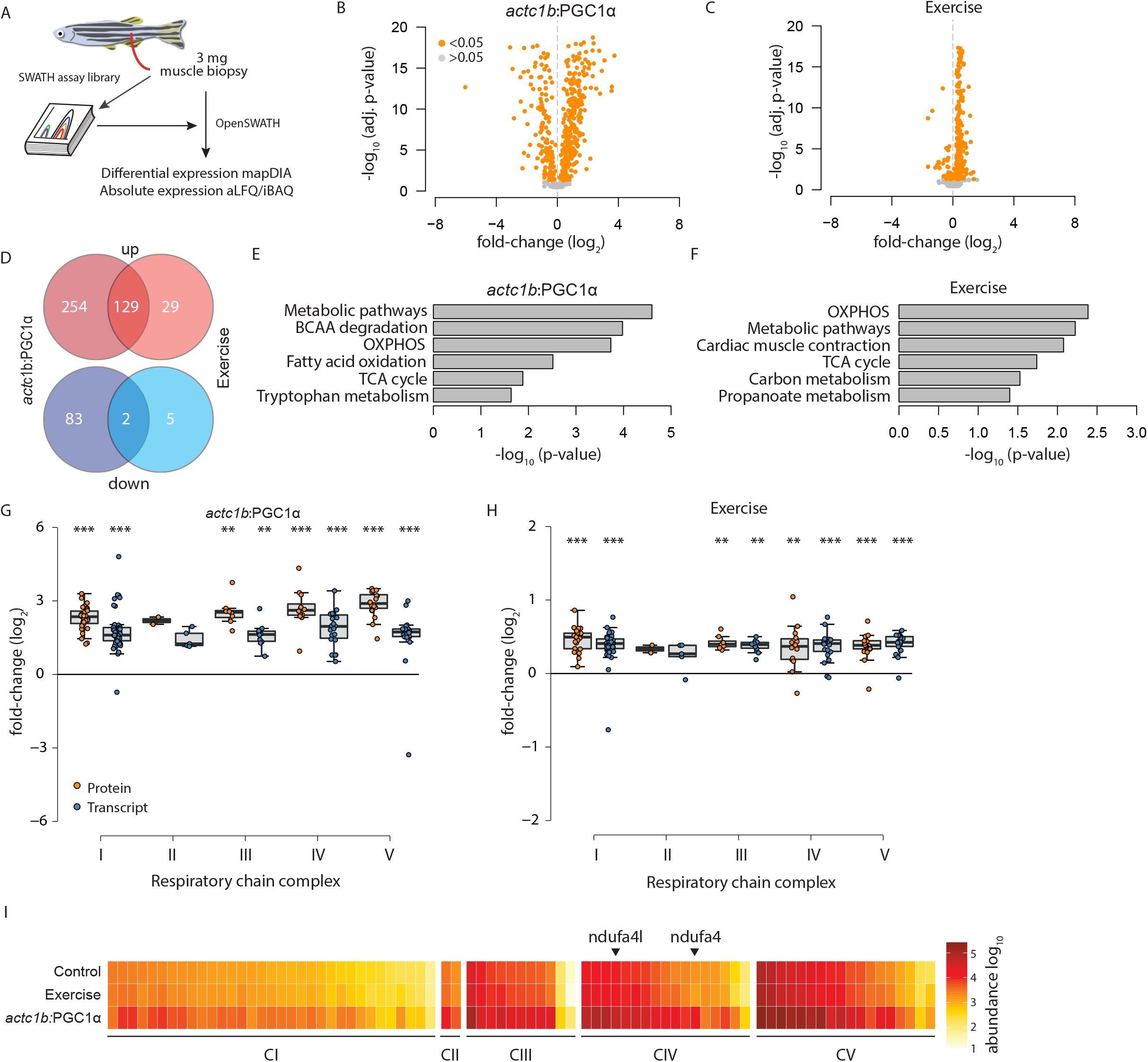
Stoichiometric induction of electron transport chain subunits in response to exercise or PGC1α. (A) Workflow of openSWATH proteomics in zebrafish. (B and C) Volcanoplot of relative protein abundance in *actc1b:*PGC1α (B) and in exercised zebrafish compared to the wild-type control animals (C) (n=8 per group). Differentially regulated proteins in orange (log2 fold-change > 0.3, p-adj. < 0.05). (D) Venn diagram of differentially regulated proteins (E and F) Gene Ontology (GO) term analysis of enriched pathways in *actc1b:*PGC1α animals (E) and in exercised animals (F) compared to controls. (G and H) Boxplots of relative protein (orange dots) and gene (blue dots) expression in *actc1b:*PGC1α animals (G) and in exercised animals (H) compared to controls. (I) Heatmap of relative protein abundance across the five complexes of the electron transport chain. ‘Abundance’ values for of ndufa4 and ndufa4l are indicated by the arrowhead. * p < 0.05, ** p < 0.01, *** p < 0.001.

To further resolve the induction of the ETC members, we analyzed the magnitude of induction of each ETC complex both for transcriptomic and proteomic changes. Indeed, for both datasets and across all complexes, PGC1α or exercise induced the majority of all members (Figure 3G and 3H). To quantify the relative abundance of the different components of ETC in skeletal muscle mitochondria, we applied the iBAQ absolute label-free quantification workflow (Rosenberger et al., 2014; Schwanhausser et al., 2011) and plotted the relative abundance of all subunits of the five OXPHOS complexes that were induced by PGC1α or exercise. Notably, using this approach we found that mitochondrial subunits belonging to complexes III to IV were higher expressed than subunits of complex I (Figure 3I). Both PGC1α overexpression and exercise did not alter the stoichiometric relationship between the different subunits.

### ndufa4l associates to free complex IV and supercomplexes

Interestingly we observed that the abudance of NDUFA4, an atypical ETC subunit first identified as a complex I member (Carroll et al., 2006), was more similar in terms of its expression pattern to CIII, CIV and CV subunits. NDUFA4 has been recently shown to co-purify with CIV was subsequently suggested as a 14^th^ subunit of Complex IV (Balsa et al., 2012). However, this view has been questioned proposing NDUFA4 to be a developmental or cancer-specific supercomplex assembly factor rather than associating to CIV in adult tissues under physiological conditions (Kadenbach, 2017). The zebrafish orthologue of human *NDUFA4* has undergone a gene duplication that resulted in two paralogues with near identical sequences: *nudfa4* and *ndufa4l* (Fig. S5). Using aLFQ/iBAQ-based quantification we found that in skeletal muscle under physiological conditions ndufa4 is expressed at very low levels, whereas ndufa4l is the major paralogue expressed (Figure S5). Comparing the abundance of ndufa4l with the abundance of the six most enriched CI or CIV members further suggests that ndufa4l is the fourth most strongly enriched CIV member expressed in skeletal muscle (Figure 4A). To confirm that ndufa4/ndufa4l is indeed associated to CIV, we performed blue native gel electrophoresis of mitochondrial crude extracts to separate ETC free complexes and supercomplexes, followed by SDS-PAGE to resolve proteins by size (2D western blot) using antibodies against ndufa4/ndufa4l or ndufs4. Indeed, in wild type conditions ndufa4/ndufa4l was detected in a pattern clearly distinct from ndufs4, a prototypical member of CI (Figure 4C). Next, we asked whether ndufa4/ndufa4l associates with CIV by co-migration with mt-co1, a prototypical member of cytochrome c oxidase. Indeed, the in-gel migration pattern of ndufa4/ndufa4l resembled that of mt-co1 (Figure 4D, left panel). To further assess the association dynamics of ndufa4l to free CIV or to CIV-containing supramolecular organizations, we compared wild-type animals to zebrafish exercised for 3 days for 3 hours per day or to *actc1b:*PGC1α transgenic zebrafish. Again, in all three conditions we found robust association of ndufa4l to CIV (Figure 4D). Remarkably, with exercise or PGC1α overexpression a shift towards increased assembly of supercomplexes was observed. The extent of this shift was similar for ndufa4l and mt-co1 (Figure 4E). Thus, ndufa4l is expressed in post-mitotic adult skeletal muscle at similar levels as the CIV core subunits and associates with CIV supercomplexes.

**Figure 4.**
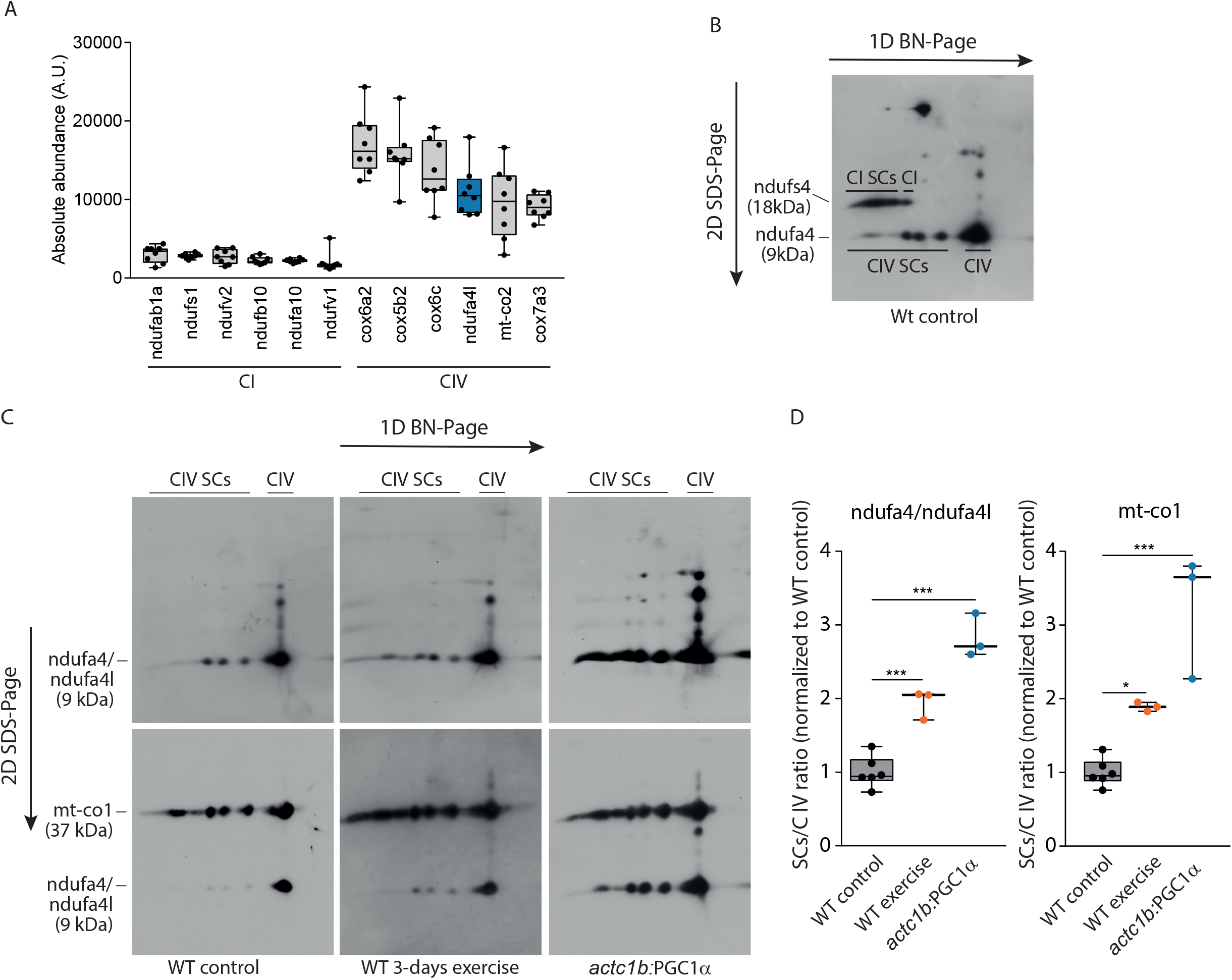
ndufa4l associates with complex IV *in vivo*. (A) Protein abundance of the six most enriched proteins of complex I and complex IV in wild-type conditions. (B) 2-D blue native/SDS-PAGE of wild-type skeletal muscle probed for ndufs4 (CI) and nudfa4l. Ndufs4 and ndufa4l show distinct migration patterns. (C) 2-D blue native/SDS-PAGE of skeletal muscle from control, 3 days exercised and *actc1b:*PGC1α animals blotted for ndufa4 (upper panels), and subsequently re-blotted for mt-co1 (lower panel). mt-co1 and ndufa4l co-migrate as free complex IV and in complex-IV associated supercomplexes. (D) Distribution of ndufa4l and mt-co1 to complex IV-containing supercomplexes relative to free complex IV. WT: wild-type, A.U.: arbitrary units. * p < 0.05, *** p < 0.001.

## DISCUSSION

Regular aerobic exercise leads to a plethora of molecular responses that translate into the many health benefits of training observed in humans. These molecular adaptations include system-wide effects on physical performance (strength, endurance) and behavior (mood, memory), suggesting that molecular signals downstream of exercise could provide novel therapeutic targets to a wide range of disease conditions (Cartee et al., 2016; Fan and Evans, 2017; Gabriel and Zierath, 2017; Handschin and Spiegelman, 2008; Penedo and Dahn, 2005). Leveraging the advantages of zebrafish, such as rapid genetics, successful generation of a large variety of disease models and the structural and metabolic similarities of skeletal muscle organization compared to human, provides a useful model to study exercise adaptations (Gut et al., 2017). The low cost and relative ease to train groups of zebrafish further facilitates utilizing zebrafish for exercise protocols. Here, we show that a relatively short protocol of four weeks of aerobic endurance training induces a molecular program that translates into stable transcriptome and proteome changes reminiscent of exercise adaptations in mammals. Furthermore, PGC1α, a key mediator of exercise effects in mammals, is sufficient to activate a molecular ‘exercise’ signature leading to increased aerobic performance, suggesting evolutionary conservation of this key pathway of physical plasticity.

In addition to these proof-of-concept demonstrations, we performed for the first time at this scale a proteomic profiling of zebrafish using SWATH-MS. Despite muscle being a particular challenging tissue to perform proteomics due to a small set of very high abundant proteins, we quantified with our PCT-SWATH-MS approach 3617 proteotypic peptides from 885 proteins across 24 biological samples from about 3mg of frozen tissue. The only previous study applying SWATH-MS in zebrafish, quantified 200 blood plasma proteins between male and female fish (Li et al., 2016a). As the SWATH-MS approach requires a spectral library, we generated a SWATH spectral library for zebrafish muscle that contains coordinates to extract data for 7989 proteotypic peptides from 1179 proteins. This represents the first SWATH spectral library provided to the community in order to support proteomic analysis of zebrafish. The proteomic results highlighted that the large majority (>80%) of the proteins induced by prolonged exercise are also significantly increased upon PGC1α overexpression. Hence, SWATH-MS proteomics is a sensitive and fast approach to profile the molecular changes induced in zebrafish tissue by specific perturbations.

The most obvious molecular changes that we observed in response to gain-of-function of PGC1α or to chronic exercise encompass concerted effects on mitochondrial bioenergetics, including the up-regulation of almost all members of the ETC on transcriptomic and proteomic levels. Mitochondrial bioenergetics, such as the capacity to metabolize carbon substrates into ATP, decline with age in skeletal muscle and contribute to the loss of muscle mass and strength, and ultimately mobility (Johnson et al., 2013; Short et al., 2005). Exercise in humans, even in aged individuals, can improve mitochondrial function and slow functional decline, which can at least partially be attributed to the improved bioenergetics capacity of skeletal muscle (Cartee et al., 2016). In this study, we describe how human PGC1α can improve the bioenergetics properties of skeletal muscle in a vertebrate model. Although the induction amplitude as well as the number of genes that become differentially regulated in response to exercise are much less pronounced compared to what can be observed in genetic gain-of-function experiments, the genome- and proteome-wide regulation of the same gene sets by both conditions suggest highly orchestrated adaptive events. This finding emphasizes the importance of assessing exercise responses at a systems level as small fold-changes of individual genes or proteins may be interpreted as false negative results, while in reality coordinated co-expression programs occur that can be detected by the appropriate statistical methods.

We show that zebrafish are capable of sustaining high swimming speeds despite a predominantly lethargic behavior in laboratory housing conditions. Furthermore they greatly improve their Ucrit within a few weeks, concomitant with a near complete enrichment of ETC subunits. This remarkable plasticity is an interesting finding as it suggests a rapid activation of cellular programs that allow resistance to fatigue. Recently it has been reported that 4 months of endurance exercise in humans induces an increase in ETC complexes as well as their reorganization into supercomplexes (Greggio et al., 2017). Here, we report that the ETC in zebrafish organizes into supercomplexes after only 3 days of exercise, and that PCG1α overexpression is sufficient to induce a shift from free to supramolecular structures. This result supports the hypothesis that ETC supercomplex formation is an early functional adaptation to exercise, although the exact role of it still remains elusive. The factors that regulate the dynamic assembly and disassembly of these supramolecular structures are to date unclear (Lenaz et al., 2010; Milenkovic et al., 2017). Our set-up in a vertebrate model can now be used to identify novel mediators of the physiological relevance of this process through genetics in conjunction with endurance training. As a proof of concept, we analyzed the relationship of ndufa4, a recently proposed supercomplex assembly factor to free and supercomplex ETC formations; NDUFA4 was originally purified as an accessory subunit of CI (Carroll et al., 2003; Carroll et al., 2006), but was recently shown to co-purify with CIV when using mild detergents, and was subsequently proposed as a 14^th^ subunit of cytochrome c oxidase (Balsa et al., 2012). Loss-of-function mutations within *NDUFA4* in human patients lead to cytochrome c oxidase deficiency and the clinical presentation of Leigh syndrome (Pitceathly et al., 2013). We found in our dataset that the translation product of *ndufa4l*, the zebrafish muscle-expressed orthologue of *NDUFA4*, is highly expressed at levels similar to the abundance of other subunits of CIV. ndufa4/ndufa4l is regulated by exercise or PGC1α maintaining a stoichiometric relationship relative to other subunits of cytochrome c oxidase, and is present in free CIV as well CIV-containing supercomplexes. Systems analyses of isogenic but across strains genetically diverse Bxd mice show that NDUFA4 is also co-expressed with members of CIV in mammals (unpublished data P.B. and R.A.). These result, together with the low affinity of ndufa4/ndufa4l for cytochrome C oxidase demonstrated by crystallization and proteomics experiments (Balsa et al., 2012; Liu et al., 2018), confirms the specific association to complex IV. The exact role of NDUFA4 in cytochrome C oxidase function or assembly remains to be elucidated.

Looking forward, the genetic tractability of zebrafish can be leveraged to iteratively test exercise performance at an elevated throughput of genetic manipulation. We have identified many genes with known functions for skeletal muscle development or physiology that have not been reported to date to be regulated by exercise. Furthermore, zebrafish allow one now to efficiently develop and test exercise paradigms that are less well explored than chronic endurance training. For example, the molecular responses to strength training are not well understood, but could potentially be studied in zebrafish using increasing levels of viscosity of the water. In summary, we expect zebrafish to become an important animal model in exercise biology.

## EXPERIMENTAL PROCEDURES

### Zebrafish husbandry and transgenic lines generation

Adult AB zebrafish were raised at 28°C under standard husbandry conditions. All experimental procedure were carried out according to the Swiss and EU ethical guidelines and were approved by the animal experimentation ethical committee of Canton of Vaud (permit VD3177). Transgenic zebrafish were generated using I-SCEI meganuclease mediated insertion into one-cell stage AB embryos of a construct harboring the zebrafish *ppargc1a* or the human *PPARGC1A* cDNA fused to a triple Flag sequence under the control of the skeletal muscle-specific *actc1b* promoter. For rapid selection of transgenic animals, the injected constructs carried an eye-marker cassette harboring *ZsGreen* under the control of the *cryaa* (alpha-crystallin A chain) promoter in reverse direction. Transgenic carriers were outcrossed with AB fish to raise transgenic and wild-type siblings.

### Endurance test

Endurance tests were performed using a 170 ml swim tunnel from Loligo^®^ Systems (#SW10000). Briefly, after a habituation phase, in which one fish was placed in the tunnel and let swim for 20 min at a low speed of 5 cm/s, the current was increased stepwise by 5 cm/s every 5 min until the fish showed fatigue. Fatigue was defined as the moment at which a fish stopped swimming and was at the rear of the swim tunnel for longer than 5 seconds. The critical swimming velocity of fish (Ucrit) was calculated as previously described by using the following equation:

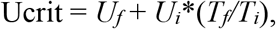

where *U_f_* is the highest velocity which could be maintained for 5 min, *U_i_* is the step increment (5 cm/s), *T_f_* the time elapsed at fatigue velocity and *T_i_* is the interval time (300 s) (Gilbert et al., 2014).

### Chronic exercise interventions

Chronic endurance swimming exercise was performed using a 5 L swim tunnel by Loligo^®^ Systems (#SW10050).

The day before the beginning of the chronic exercise intervention, 8 *actc1b:*PGC1α and 16 wild-type siblings (4 months of age) were measured in body size and weight and were subjected to an endurance test in the 5 L swim tunnel to determine their exercise speed. Baseline Ucrit was determined as the speed by which the weakest zebrafish fatigued (5 seconds at the rear of the tunnel). Based on this result, 8 wild-type fish were trained for 4 consecutive weeks, 5 days per week, 3 hours per day at increasing swimming speeds: 3 days at 50 cm/s (75% of baseline Ucrit), 5 days at 55 cm/s, 5 days at 60 cm/s, 5 days at 65 cm/s, 4 days at 70 cm/s. As control group, the 8 *actc1b*:PGC1α and the remaining 8 wild-type fish were placed in the swim tunnel for 3h per day at a low swim speed of 5 cm/s. Two days before the end of the intervention individual endurance tests were performed on all the three groups. At the end of the 4 weeks, fish were euthanized after the last training session and trunk muscle was isolated, flash frozen in liquid nitrogen and processed for RNA and protein extraction.

A second chronic exercise intervention with an identical design was conducted on another cohort (n=5 zebrafish for each of the three conditions) in order to perform immunofluorescent analysis of morphometry and mitochondria content (Figure 2C and D).

For the short exercise protocol (Figure 4), fish were exercised for three consecutive days, 3 hours per day, at 75% of their Ucrit. The zebrafish were euthanized for 2D western blot analysis of supercomplex formation in the morning of the next day.

### Immunohistology and microscopy

Skeletal muscle were processed for freezing, cryosectioning and immunofluorescent staining as previously described (Parisi et al., 2015). Primary antibodies: anti-Laminin (Sigma, #L9393), anti-Slow Myosin Heavy Chain (DSHB, #BA-D5), anti-Tom20 (SCBT, #sc-11415). Secondary Antibodies: Alexa Fluor 488 Goat anti-Mouse IgG2b (Thermo Scientific, #A-21141), Alexa Fluor 555 Goat anti-Rabbit IgG (Thermo Scientific, #A-21428). Microphotographs were acquired with a Leica DMI Inverted Microscope 4000B and a Leica DMC2900. Fluorescence intensity and myofibers cross sectional area were quantified by using the ImageJ software.

### Gene expression and RNA-seq

For gene expression analysis, flash frozen skeletal muscle was lysed in Qiazol with the FastPrep®-24 tissue homogenizer (MP-Biomedials). Total mRNA was extracted using the QIAcube plateform and mRNAeasy kit (Qiagen).

For RT-qPCR, cDNA was synthesized using standard cDNA synthesis kit following the manufacturer’s instructions (Thermo Fisher). qPCR was performed with the Roche Light Cycler 480 using SYBR green kit (Maxima SYBR Green, Thermo Fisher) and the following primers: *βactin-Fwd*: GTGGTCTCGTGGATACCGCAA, *βactin-Rev*: CTATGAGCTGCCTGACGGTCA, *ppargc1a-Fwd*: GCGAGGGAACGAGTGGATTT, *ppargc1a-Rev*: CTCTCCACACCGAATCCTGA, *3xFlag-Fwd:* GATGTCGGACCACACGGAC, *PPARGC1A-Rev*: TGGACTACAAAGACCATGACGG.

Microarrays on wild-type and *actc1b*:PGC1α skeletal muscle of 4-months old zebrafish were performed by the OakLabs company (http://www.oak-labs.com/). After quality control with the 2100 Bioanalyzer using the RNA 6000 Pico Kit, RNA was labelled using the Low Input QuickAmp Labeling Kit (Agilent Technologies) and cRNA labelled with cyanine 3-CTP is obtained by in vitro transcription. For hybridization, the Agilent Gene Expression Hybridization Kit (Agilent Technologies) is used and 600 ng of each cRNA are hybridized on 8×60K microarrays at 65 C for 17 h using Agilent’s recommended hybridization chamber and oven. Microarrays were then washed following the Agilent Gene Expression protocol and scanned using the SureScan Microarray Scanner (Agilent Technologies). From the raw data, the control probes were removed and the signal across replicates and probes of the same target were averaged. Data were then quantile normalized and genes were tested for differential expression in wild-type and *actc1b*:PGC1α condition via the moderated t-statistic as implemented in Limma (Ritchie et al., 2015). Genes differentially expressed were associated to a list of known genes that are regulated by chronic exercise and t-stat values were represented as boxplots by pathway. Statistical analysis was performed with a gene set enrichment analysis (Subramanian et al., 2005) on t-stats with Benjamini-Hochberg multiple test correction.

For RNA-seq, RNA quantification was performed with Ribogreen (Life Technologies) and quality was assessed on a Fragment Analyzer (Advances Analytical). Sequencing libraries were prepared from 250 ng RNA using the TruSeq Stranded mRNA LT Sample Prep Kit (Illumina) following the manufacturer’s protocol, except for the PCR amplification step. The latter was run for 15 cycles with the KAPA HiFi HotStart ReadyMix (Kapa BioSystems). This optimal PCR cycle number has been evaluated using the Cycler Correction Factor method as previously described (Atger et al., 2015). Libraries were quantified with Picogreen (Life Technologies). The size pattern was controlled with the DNA High Sensitivity Reagent kit on a LabChip GX (Perkin Elmer). Libraries were pooled and the pool was clustered at a concentration of 9 pmol on 2*8 lanes of paired-end sequencing high output flow cell (Illumina). Sequencing was performed for 2 x 125 cycles on a HiSeq 2500 with v4 SBS chemistry following Illumina’s recommendations.

### RNA-seq data analysis

Image analysis and base calling were performed using the Illumina Real-Time Analysis. Raw data are available at (accession number #). Paired-end reads were mapped on the reference genome of zebrafish GRCz10 using STAR 2.4.0i (Dobin et al., 2013). Uniquely mapped reads were counted for each gene using the Python Package HTseq 0.9.1 (Anders et al., 2015) to determine the expression level of transcripts. Normalization of the read counts and differential expression analysis were performed using the Bioconductor 3.6 Package DESeq2 (Love et al., 2014). Genes with adjusted p-values smaller than 0.05 and log2 fold-changes larger than 0.5 were used to compare genotype- and exercise-induced expression.

Gene ontology (GO) analysis was performed using DAVID Bioinformatics Resources 6.8 (Dennis et al., 2003). KEGG pathway annotation was used for the enrichment test. Categories with p-value smaller than 0.05 were considered as significantly enriched. A ranked gene set enrichment analysis was performed to clarify the correlation between PGC1α overexpression and the gene signature of exercise. Gene set was composed by genes that were significantly regulated in exercised zebrafish (p-adj. <0.05, log2 fold-change >1). Barcode-plot were used to display the enrichment of the gene set in the ranked gene list. The same analysis was used to show the enrichment of oxidative phosphorylation pathway using a list of gene selected from KEGG annotation.

### Total protein isolation and western blotting

Total proteins were extracted from flash frozen skeletal muscle with RIPA Buffer (Sigma) supplemented with Protease Inhibitor Cocktail (Thermo Fisher Scientific) using a tissue Homogenizer. Protein quantification was performed using the BCA assay kit (Thermo Fisher Scientific). 20 μg of proteins were loaded onto precast Novex gels (Thermo Fisher Scientific) and transferred onto nitrocellulose membranes using the iBlot2 instrument (Thermo Fisher Scientific). Primary Antibodies: Porin (Abcam, #ab15895), Tubulin (Abcam, #ab6046), Ndufa4 (Sigma, #SAB4501963), Ndufs4 (Thermo Fisher Scientific, #PA5-21677), Mt-Co1 (Abcam, #ab14705). Secondary Antibodies: anti-rabbit IgG HRP-labeled (Perkin Elmer, #NEF812001EA), anti-mouse IgG HRP-labeled (Perkin Elmer, #NEF822001EA)

### Protein expression and Mass Spectrometry

For protein expression analysis by mass spectrometry, flash frozen skeletal muscle was homogenized by manual grinding while keeping the samples frozen with liquid nitrogen. From the resulting powder, 3 mg were transferred into the pressure-cycling technology (PCT) Microtubes for subsequent lysis. The lysis and digestion was performed with the Barocycler NEP2320EXT (Pressure BioSciences) at 31°C based on a protocol described previously (Guo et al. 2015). The tissue was lysed using lysis buffer (6 M urea, 2 M thiourea, 100 mM ammonium bicarbonate, 5 mM EDTA, cOmplete protease inhibitor (1:50), pH 8.0) and applying 60 cycles (50 s at 45 kpsi followed by 10 s at atmospheric pressure) in microtubes containing a MicroPestle. For the subsequent reduction and alkylation, the urea was diluted to 3.75 M with 100 mM ammonium bicarbonate and the samples were incubated for 30 min in the dark at 25°C with 9 mM tris(2-carboxyethyl)phosphine (TCEP) and 35 mM iodoacetamide. The proteins were then first digested with LysC (enzyme/protein ratio of 1:100) and 45 cycles (50s at 20 kpsi followed by 10s at atmospheric pressure). After diluting the urea with 100 mM ammonium bicarbonate to 2 M, the samples were digested a second time with trypsin (enzyme/protein ratio of 1:75) and 90 cycles (50 s at 20 kpsi followed by 10 s at atmospheric pressure). The digestion was stopped by acidifying the samples with trifluoroacetic acid to pH 2 and the digested peptides were then desalted on C18-columns (The Nest Group Inc.). On the columns, the peptides were washed with 2% acetonitrile and 0.1% trifluoroacetic acid in H_2_O, eluted with 50% acetonitrile and 0.1% trifluoroacetic acid in H_2_O and subsequently dried in a centrifugal vacuum concentrator. Before injection to the mass spectrometer, the dried peptides were dissolved in 2% acetonitrile and 0.1 formic acid in H_2_O and iRT peptides (Biognosys) were added to the sample.

To measure the samples by mass spectrometry, 2μg peptides were first separated by nano-flow liquid chromatography (NanoLC Ultra 2D, Eksigent) and then quantified on an ABSciex TripleTOF 5600 instrument. The reverse-phase chromatography was performed on a fused silica PicoTip™ Emitter (inner diameter 75 mm) (New Objective, Woburn, USA) packed with C18 beads (MAGIC, 3 μm, 200 Å, BISCHOFF, Leonberg, Germany) for 21 cm. The peptides were separated with a flow of 300 nl/min and a linear 90 min gradient from 2% to 35% Buffer B (98% acetonitrile and 0.1% formic acid in H_2_O) in Buffer A (2% acetonitrile and 0.1% formic acid in H_2_O). After ionization, the precursor peptide ions were accumulated for 250ms with 64 overlapping variable mass-to-charge windows between 400 Th and 1200 Th. Fragmentation of the precursor peptides was achieved by Collision induced dissociation (CID) with rolling collision energy for peptides with charge 2+ adding a spread of 15eV. The MS2 spectra were acquired in high-sensitivity mode with an accumulation time of 50 ms per isolation window resulting in a cycle time of 3.5 seconds. Samples and controls from the biological replicates were injected in a block design.

### Mass Spectrometry data analysis

The quantitative data was extracted from the SWATH-MS mass spectra using the OpenSWATH workflow (Röst et al., 2014) on the in-house iPortal platform (Kunszt et al., 2015) using a zebrafish muscle SWATH spectral library (see below). An m/z fragment ion extraction window of 0.05 Th, an extraction window of 600 s, and a set of 10 different scores were used as described before (Röst et al., 2014). To match features between runs, detected features were aligned using a spline regression with a target assay FDR of 0.01 (Röst et al., 2016). The aligned peaks were allowed to be within 3 standard deviations or 60 s after retention time alignment. For runs where no peak could be identified, the signal was integrated using the single shortest Path method (Rost et al. 2016). The data was then further processed using the R/Bioconductor package SWATH2stats (Blattmann et al., 2016) where precursor had to pass an m-score threshold of 2.5E-05 in at least 20% of the 24 files to be selected for further analysis. This stringent threshold resulted in an estimated precursor FDR of 0.0017, peptide FDR of 0.0021, and protein FDR of 0.0097 (using an estimated fraction of false targets (FFT) or π0-value of 0.48). A total of 4227 proteotypic peptides from 905 proteins passed this stringent threshold.

For differential expression analysis, mapDIA v3.0.2 (Teo et al. 2015) was used. For proteins where many peptides were quantified, the 10 peptides with the highest signal were selected. Furthermore, mapDIA was used with the following settings: Local total intensity normalization within a retention time window of 10 min; An independent study design; Minimum correlation of 0.25; Standard deviation factor of 2; Between 3 and 5 fragments per peptide; At least one peptide per protein. This approach resulted in 3617 proteotypic peptides from 885 proteins being quantified by mapDIA. The absolute label-free protein abundance estimation was performed with the R/CRAN aLFQ package (Rosenberger et al., 2014) using the iBAQ method (Schwanhausser et al., 2011), selecting all peptides from the data obtained after filtering with SWATH2stats.

The heatmaps depicting the estimated absolute abundance for proteins in the respiratory chain complexes contain intensities that were log10-transformed and averaged for each biological replicate. For the other heatmaps and boxplots, the mapDIA results were used. Both heatmaps and boxplots were generated using the R/CRAN ggplot2 package.

### Generation of SWATH spectral library

To generate the SWATH spectral library, in total 42 samples from zebrafish muscle were utilized consisting of 12 unfractionated samples prepared as described above, 24 samples for which the peptides were separated using off-gel isoelectric focusing into 12 fractions, and 6 samples for which mitochondria were enriched using the protocol from Frezza et al. 2007 (Frezza et al., 2007). Off-gel-fractionation was performed on an Agilent 3100 OFFGEL Fractionator according to the manufacturer’s protocol. The 42 samples were measured by injecting them once in data-dependent acquisition (DDA) mode on an ABSciex TripleTOF 5600 instrument (Sciex, Concord, Canada). The peptides were quantified as described above using a 120min, instead of a 90min, gradient and operating the machine in DDA mode. Precursor selection on the MS1 level was performed with a Top20 method using an accumulation time of 250 ms and a dynamic exclusion time of 20 s. The MS1 spectra were obtained in an m/z range from 360 to 1460. For MS2 spectra, only fragments with a charge state from 2 to 5 were selected using an accumulation time of 150 ms. The spectral library was built using an in-house computational pipeline configured on the iPortal platform (Kunszt et al., 2015). Raw files from the DDA runs were converted into centroided mzXML files using ProteoWizard version 3.0.8851, searched using X!Tandem and Comet and the peptides passing an mayu-protFDR of 0.01 were incorporated into a spectral library using the workflow outlined previously (Schubert et al., 2015). As a protein sequence database, the Ensembl 87 release for Danio rerio was used (Danio_rerio.GRCz10.pep.all.fa.gz from dec2016.archive.ensembl.org) selecting only the proteins whose transcripts were detected in the transcriptome of muscle tissue from wild type and actc1b:PGC1α fishes. To these proteins, the 13 mitochondrial proteins were added (mt-nd1, mt-nd2, mt-nd3, mt-nd4, mt-nd4l, mt-nd5, mt-nd6, mt-co1, mt-co2, mt-co3, mt-atp8, mt-atp6, and mt-cyb).

### Data availability

The mass spectrometry proteomics data have been deposited to the ProteomeXchange Consortium via the PRIDE partner (Vizcaino et al., 2016) repository with the dataset identifier ProteomeXchange: PXD009811 (Reviewer account details: Username. Password: isgb24vf)

### Mitochondria isolation and 2DS Blue Native/SDS-PAGE

Mitochondria crude extracts were prepared from fish trunk muscles as previously described, with minor changes (Fernandez-Vizarra et al., 2002). After BCA quantification of protein concentration, samples were incubated 1 hour at 4°C with 4% Digitonin. Non solubilized fraction was pelleted by centrifugation and discarded, while the supernatant containing mitochondria extracts was mixed with Native Page Buffer and stored at -80°C. 25 μg of mitochondrial crude extracts were loaded onto NativePAGE™ 3-12% Bis-Tris Protein Gels (Thermo Fisher Scientific) and gel migration was performed following manufacturer’s instructions.

At the end of the migration, gel lanes were cut individually and incubated 1 hour in MES buffer supplemented with 1% SDS 1% beta-mercaptoethanol. Each lane was inserted into NuPAGE™ 4-12% Bis-Tris Protein Gels, 1.0 mm, 2D-well and let migrate as a normal SDS-PAGE gels.

Proteins were then transferred onto nitrocellulose membranes and the protocol for regular western blot was followed (see above).

### ETC complex activity assay

The specific activity of each mitochondrial complex and of the citrate synthase has been measured by a spectrophotometric assay described in (Bugiani et al., 2004).

### Statistical analysis

All numerical data are expressed as mean ± SEM and reported as histograms or boxplots. Student’s unpaired two-tail t test and One-Way Anova test were used for statistical analysis unless otherwise stated. Differences were considered statistically significant for p < 0.05.

## Supporting information

## AUTHOR CONTRIBUTIONS

P.G. and B.S. conceived the project. P.B., V.S. established the SWATH-MS workflow and analysis. A.P. and G.L. performed zebrafish experiments. E.M., B.W. and C.G. helped with bioinformatics analyses. L.S., J.R., G.C. contributed experiments. A.C. and F.R. performed RNA processing and RNAseq experiments. P.G., R.A. and B.S. supervised the work. P.G., A.P., P.B. and B.S. wrote the paper. All authors read and commented on the paper.

## ACKNOWLEDGEMENTS

We thank Thierry Guillaud for excellent zebrafish husbandry and Sebastien Cotting for technical facility support. This study was supported by grant TPdF 2013/134 of the Swiss SystemsX.ch initiative evaluated by the Swiss National Science Foundation to P.B. as well as grant NIH DK 61562 to B.M. The R.A. group is supported by the Swiss National Science Foundation (grant no. 3100A0-688 107679), the European Research Council (ERC-2014-AdG 670821), ETH Zurich, and SystemsX.ch. The Nestlé Institute of Health Sciences is member of the Lausanne Integrative Metabolism & Nutrition Alliance.

---

**Figure S1. Alignment of zebrafish and human PGC1A protein sequence**

Zebrafish and human PGC1A protein sequences show 61.45% identity. Alignment was performed with the Clustal Omega tool (https://www.ebi.ac.uk/Tools/msa/clustalo/). All the functional and regulatory domains are highly conserved between the 2 species. These domains include: the LxxLL motifs (PPAR/NR interaction domains) in pink, the phospho-Threonine 178 and phospho-Serine 539 in red, the phosphodegron motifs in yellow, the arginine/serine rich domains in green and the RNA recognition motif in blue.

**Figure S2. Human PGC1α induces morphological and metabolic reprogramming at 6 weeks**

(A and B) Lateral view of trunks of a wild-type (A) and *actc1b:*PGC1α (B) zebrafish at 6 weeks of age. (C and D) Immunofluorescence staining of slow-myosin heavy chain as a marker for slow-twitch myofibers and laminin as a marker for the basal lamina of myofibers in trunk muscle of wild-type (C) and *actc1b:*PGC1α zebrafish (D) (Scale bar, 100μm). (E) Average cross-sectional area of trunk muscle fibers in wild-type and transgenic animals (average counts from at least 250 myofibers per animal, n=6 per genotype). (F) Frequency distribution of cross-sectional fiber area in wild-type and PGC1α zebrafish (counts from at least 250 myofibers per animal, n=6 per genotype). No significant difference was detected between fiber area distribution of wild-type and PGC1α animals. KS test. ** p<0.01

**Figure S3. Increased citrate synthase and ETC complex specific activity upon PGC1A overexpression**

Quantification of the specific enzymatic activity of citrate synthase (A), Complex I (B), Succinate Dehydrogenase (C), Complex II (D) and Complex IV (E). n=5 per genotype. *** p<0.001; **** p<0.0001

**Figure S4. Enrichment analysis of gene ontology terms of gene signatures in response to PGC1α and exercise**

(A and B) Gene Ontology (GO) term analysis of enriched pathways in *actc1b:*PGC1α animals (A) and in exercised animals (B) compared to controls. (C and D) Enrichment test for gene set related to ‘oxidative phosphorylation’ in *actc1b*:PGC1α animals (C) and in exercise zebrafish (D) compared to controls.

**Figure S5. ndufa4 and ndufa4l alignment and expression level**

(A) Amino acid sequence alignment of ndufa4 and ndufa4l (B) Absolute expression of ndufa4 and ndufa4l in zebrafish skeletal muscle.

**Table S1 Curated list of ‘exercise’ genes used for sub-analysis of microarray experiments presented in Figure 1**

**Table S2 Genes differentially regulated in response to Exercise and PGC1α overexpression**

**Table S3 Proteins differentially regulated in response to Exercise and PGC1α overexpression**

