## Supplementary material for "PGC1α and Exercise Adaptations in Zebrafish"

```
Zebrafish      MAWDRCNQD--SVWRELECAALVGEDQPLCPDLPELDLSELDVSDLDADSFLGGLKWYSD  58
Human          MAWDMCNQDSESVWSDIECAALVGEDQPLCPDLPELDLSELDVNDLDTDSFLGGLKWCSD  60
               **** ****  *** ::**************************.***:********* **

Zebrafish      QSEIISSQYGNEASNLFEKIDEENEANLLAVLTETLDSIPVDEDGLPSFEALADGDVTNA  118
Human          QSEIISNQYNNEPSNIFEKIDEENEANLLAVLTETLDSLPVDEDGLPSFDALTDGDVTTD  120
               ******.**.** **:********************:**********:**:*****.

Zebrafish      SDQSCPSTPDGSPRTPEPEEPSLLKKLLLAPANSQLSYNQYPGGKAQNHAASNQRIRPAP  178
Human          NEASPSSMPDGTPPPQEAEEPSLLKKLLLAPANTQLSYNECSGLSTQNHANHNHRIRTNP  180
               .: *  * ***:*   * ***************:*****:  *  .:****  *:***  *

Zebrafish      AVAKTENPWNSKPRGACPNR-SMRRPCTELLKYLTSSDEAFQTKAGEAKSTWTGCGKDRG  237
Human          AIVKTENSWSNKAKSICQQQKPQRRPCSELLKYLTTNDDPPHTKPTENRNSS--------  232
               *:.**** *..* :. * ::   ****:*******:.*:  :**  * :.:

Zebrafish      GACISSCSSSSSPSSSSTSSFSSLSSCSSSTASKKKTSSASPSSQQQQLAVQAQRAKPTI  297
Human          -----------------------RDKCTSK----KK--------SHTQSQSQHLQAKPTT  257
                                      ..*:*.    **        .: *   *  :****

Zebrafish      LPLPLTPESPNDHKGSPFENKTIERTLSVEICGTPGLTPPTTPPHKASQENPFKVSLKNK  357
Human          LSLPLTPESPNDPKGSPFENKTIERTLSVELSGTAGLTPPTTPPHKANQDNPFRASPKLK  317
               * ********** ***************:.** ************.*:***:.* * *

Zebrafish      LSSCSPSALTSKRPRLSNGGSCPQPTSGSIRKGPEQTELYAQLSKASSTMPQGGLEDRRG  417
Human          SSCKTVVPPPSKKPRYSESSG--TQGNNSTKKGPEQSELYAQLSKSS--VLTGGHEERKT  373
                *. :     **:** *:...     ..* :*****:********:*  :  ** *:*:

Zebrafish      KRPMPRVFGDHDYCQSTSTKRDSTTPAAVVPGPTEGRHVECKDLNMPTSTTTTSSLSSTP  477
Human          KRPSLRLFGDHDYCQSINSKTEILINIS--QELQDSRQLENKDVSSDWQG----------  421
               ***  *:********* .:* :     :       :.*::* **:.   .

Zebrafish      PSSSSLARQLQGLSPTPQEACPDTYAHVQHHDSSSKMTMDCSSGGRKLLRDQEIRDELNK  537
Human          --------QI--CSSTDSDQC---YL--RETLEASKQVSP--CSTRKQLQDQEIRAELNK  464
                       *:   * * .: *   *   :.  .:** .    .. ** *:***** ****

Zebrafish      HFGKPQQAFYSGVVGEPRGKQPIEDSDSGDEYPGLLGDYIHPGLPD---FEDLEVGRERL  594
Human          HFGHPSQAVFDDEADKTG---ELRDSDFSNEQFSKLPMFINSGLAMDGLFDDSEDESDKL  521
               ***:*.**.:.. ..:      :.*** .:*  . *  :*: **     *:* *   ::*

Zebrafish      FYLGEGSPLELLLEGSPSSSPSSSSFSWCSVSPPSSQLSPQHLRWPRSISRSRSRSSSHH  654
Human          SYPWDGTQSYSLFNVSPSC----SSFN----SPCRDSVSPPKSLFSQRPQRMRSRSRSFS  573
                *  :*:    *:: ***.    ***.    **  ..:** :  : :  .* **** *.

Zebrafish      RRRSLSRSPYSRSGSPS--SRSPSWSPRNMDESTFTPRICG----NPQSQSHSLFGRRPR  708
Human          RHRSCSRSPYSRSRSRSPGSRSSSRSCYYYESSHYRHRTHRNSPLYVRSRSRSPYSRRPR  633
               *:** ******** * *  *** * *    :.* :  *          :*:*:* :.****

Zebrafish      YDSYEEYQHERLKREEYRRDYEKRECERAEQRERQRQKAIEERRVVYVGRLRADSTRTEL  768
Human          YDSYEEYQHERLKREEYRREYEKRESERAKQRERQRQKAIEERRVIYVGKIRPDTTRTEL  693
               *******************:*****.***:***************:***::* *:*****

Zebrafish      KRRFEVFGEIEESTVNLRHDGDNFGFITYRYTCDALAALENGHTLRRSNEPHFELCLGGQ  828
Human          RDRFEVFGEIEECTVNLRDDGDSYGFITYRYTCDAFAALENGYTLRRSNETDFELYFCGR  753
               : **********.*****.***.:***********:******:*******  .*** : *:

Zebrafish      KQYSKSNYTDLDSHSDDFDPASTKSKYDSMDFDSLLREAQYSLRR      873
Human          KQFFKSNYADLDSNSDDFDPASTKSKYDSLDFDSLLKEAQRSLRR      798
               **: ****:****:***************:******:*** ****
```

### Supplementary Figure 1

**Figure S1. Alignment of zebrafish and human PGC1α protein sequences**

Zebrafish and human PGC1A protein sequences show 61.45% identity. Alignment was performed with the Clustal Omega tool (https://www.ebi.ac.uk/Tools/msa/clustalo/). All the functional and regulatory domains are highly conserved between the 2 species. These domains include: the LxxLL motifs (PPAR/NR interaction domains) in pink, the phospho-Threonine 178 and phospho-Serine 539 in red, the phosphodegron motifs in yellow, the arginine/serine rich domains in green and the RNA recognition motif in blue.

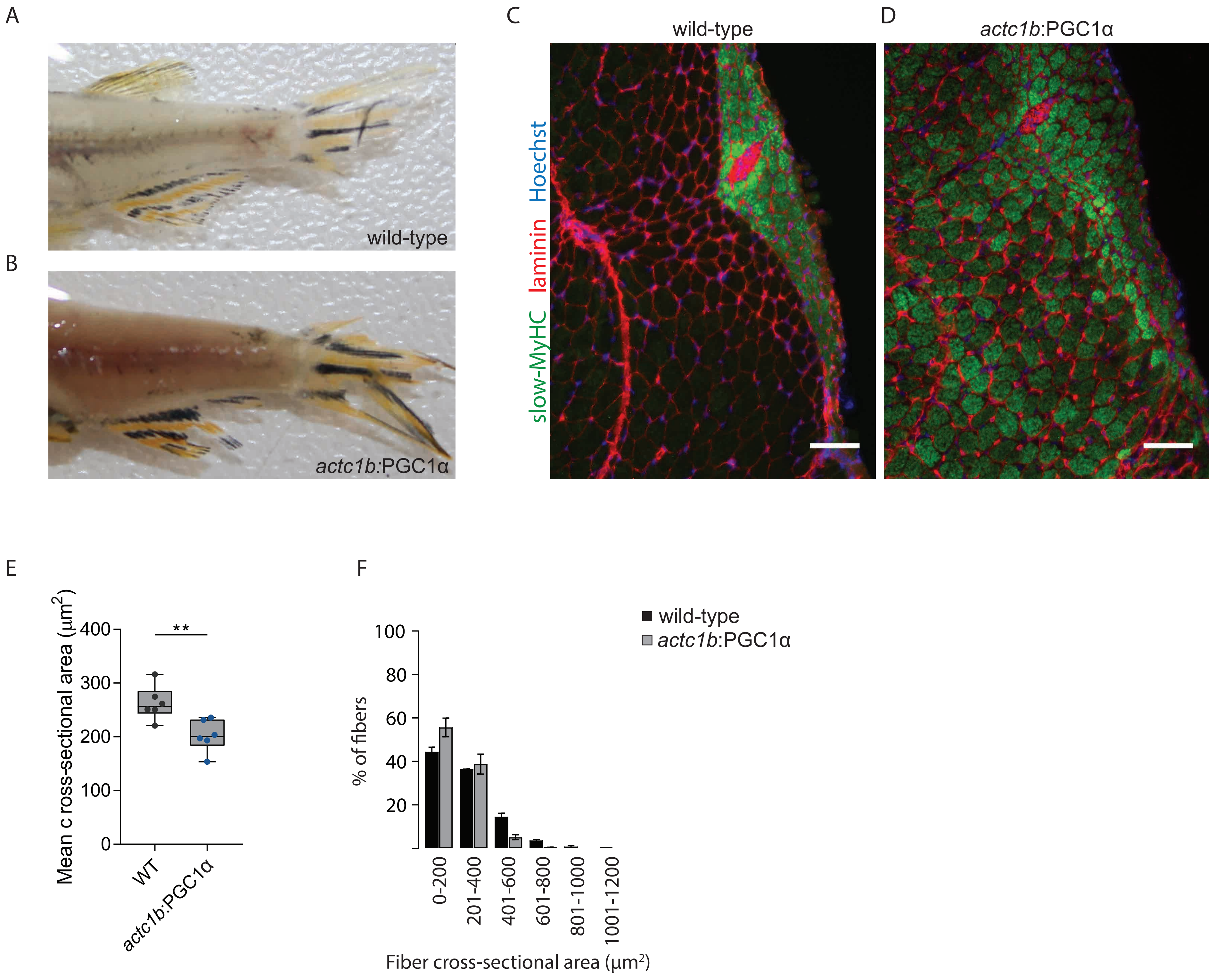


Supplementary Figure 2

**Figure S2. Human PGC1α induces morphological and metabolic reprogramming at 6 weeks**

(A and B) Lateral view of trunks of a wild-type (A) and *actc1b:*PGC1α (B) zebrafish at 6 weeks of age. (C and D) Immunofluorescence staining of slow-myosin heavy chain as a marker for slow-twitch myofibers and laminin as a marker for the basal lamina of myofibers in trunk muscle of wild-type (C) and *actc1b:*PGC1α zebrafish (D) (Scale bar, 100μm). (E) Average cross-sectional area of trunk muscle fibers in wild-type and transgenic animals (average counts from at least 250 myofibers per animal, n=6 per genotype). (F) Frequency distribution of cross-sectional fiber area in wild-type and PGC1α zebrafish (counts from at least 250 myofibers per animal, n=6 per genotype). No significant difference was detected between fiber area distribution of wild-type and PGC1α animals. KS test. ** $p<0.01$

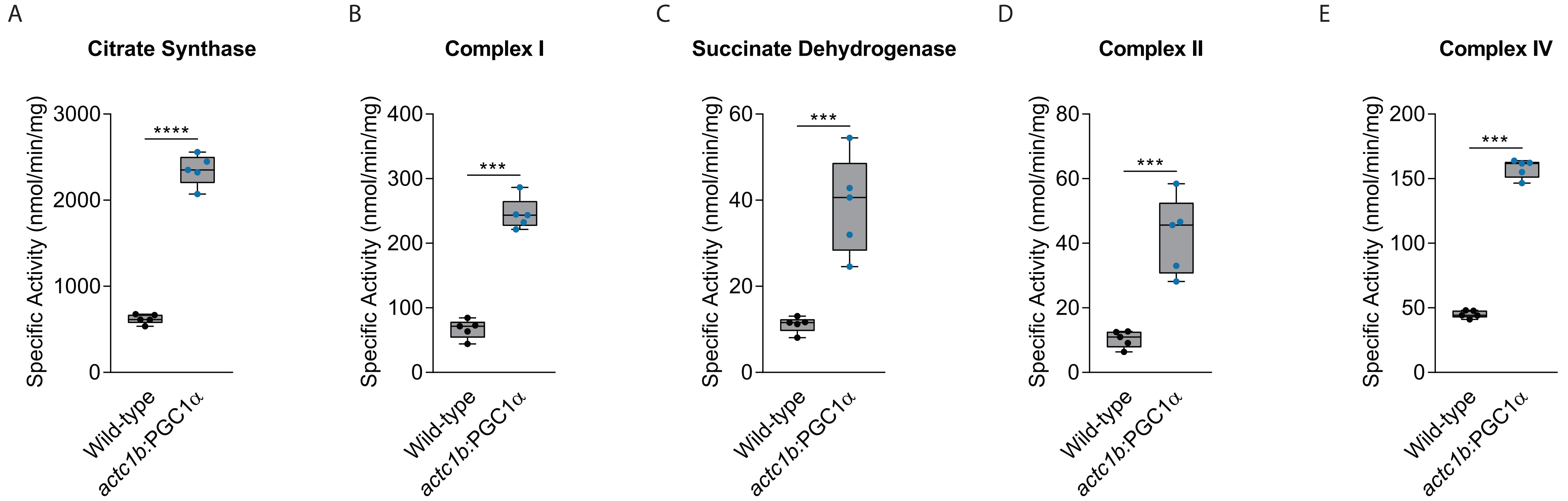


Supplementary Figure 3

**Figure S3. Increased citrate synthase and ETC complex specific activity upon PGC1A overexpression**

Quantification of the specific enzymatic activity of citrate synthase (A), Complex I (B), Succinate Dehydrogenase (C), Complex II (D) and Complex IV (E). n=5 per genotype. *** $p<0.001$; **** $p<0.0001$

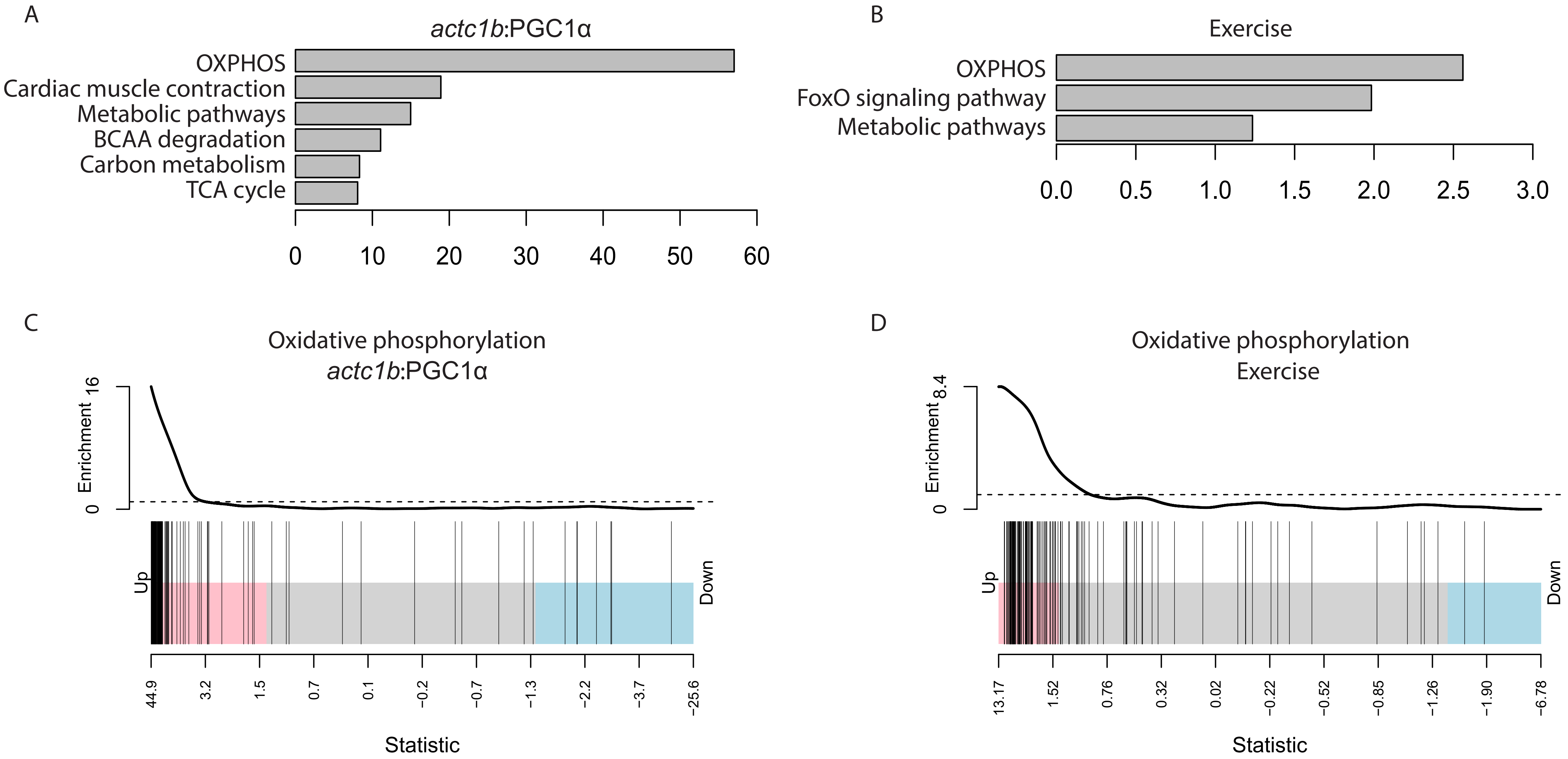


Supplementary Figure 4

**Figure S4. Enrichment analysis of gene ontology terms of gene signatures in response to PGC1α and exercise**

(A and B) Gene Ontology (GO) term analysis of enriched pathways in *actc1b:*PGC1α animals (A) and in exercised animals (B) compared to controls. (C and D) Enrichment test for gene set related to ‘oxidative phosphorylation’ in *actc1b*:PGC1α animals (C) and in exercise zebrafish (D) compared to controls.

A

```
ndufa4   MLATVMKQLKSHPALIPLFIFIGGGATMSMLYLGRLALKNPDCSWDRKNNPEPWNKLGPNDQYKLFSVNMDYSKLKKDRPDF
ndufa4l  MLSMVSRQLRSHPALIPLFIFIGGGCTMSLSYLARLALRNPDVCWDKKNNPEPWNKMGPTDQYKFYAVNMDYSKLKKNGPDF
         **: * :**:***************.***: **.****:*** .**:*********:**.****:::**********: ***
```

B

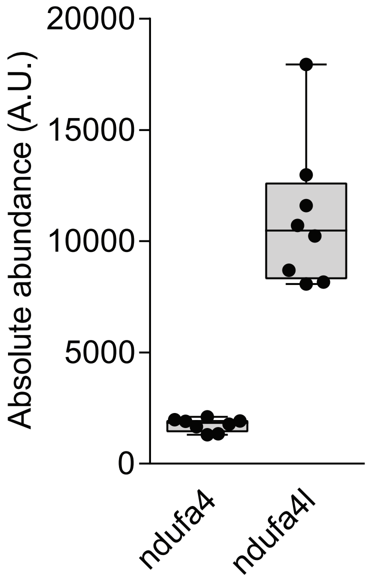


Supplementary Figure 5

**Figure S5. ndufa4 and ndufa4l alignment and expression level**

(A) Amino acid sequence alignment of ndufa4 and ndufa4l (B) Absolute expression of ndufa4 and ndufa4l in zebrafish skeletal muscle.
